# Visualization of intracellular ATP dynamics in the whole kidney under pathophysiological conditions using the kidney slice culture system

**DOI:** 10.1101/2024.05.11.592491

**Authors:** Shigenori Yamamoto, Shinya Yamamoto, Masahiro Takahashi, Akiko Mii, Akihiro Okubo, Naoya Toriu, Shunsaku Nakagawa, Takaaki Abe, Shingo Fukuma, Hiromi Imamura, Masamichi Yamamoto, Motoko Yanagita

## Abstract

ATP depletion plays a central role in the pathogenesis of kidney diseases. Recently, we reported spatiotemporal intracellular ATP dynamics during ischemia reperfusion (IR) using GO-ATeam2 mice systemically expressing an ATP biosensor. However, observation from the kidney surface did not allow visualization of deeper nephrons or accurate evaluation of ATP synthesis pathways. Here we established a novel ATP imaging system using slice culture of GO-ATeam2 mouse kidneys, evaluated ATP synthesis pathway, and analyzed intracellular ATP dynamics using *ex vivo* IR and cisplatin nephropathy models. We demonstrated that proximal tubules (PTs) were strongly dependent on oxidative phosphorylation (OXPHOS), whereas podocytes relied on both OXPHOS and glycolysis using ATP synthesis inhibitors. We also confirmed that the *ex vivo* IR model could recapitulate ATP dynamics *in vivo*; ATP recovery in PTs after reperfusion varied depending on ischemia length, whereas ATP in distal tubules (DTs) recovered well even after long ischemia. After cisplatin administration, ATP levels in PTs decreased first, followed by a decrease in DTs. An organic cation transporter 2 inhibitor suppressed cisplatin uptake in kidney slices, leading to better ATP recovery in PTs, but not in DTs. Finally, we confirmed that a mitochondria protection reagent delayed cisplatin-induced ATP decrease in PTs. This novel system may provide new insights into the energy dynamics and pathogenesis of kidney disease.

## Introduction

The kidney consumes a large amount of adenosine-5ʹ-triphosphate (ATP) for active reabsorption and glycogenesis in the tubules as well as for maintaining the structural integrity of glomerular filtration. It is one of the most ATP-consuming organs, along with the heart.^1^ ATP is produced by two major pathways: oxidative phosphorylation (OXPHOS) and glycolysis. Mitochondria are important intracellular organelles responsible for ATP synthesis through OXPHOS.^2,3^ In addition, mitochondrial damage not only causes energy depletion but also results in renal fibrosis through an inflammatory response^4,5^ and has been reported in common diseases such as acute kidney injury, diabetic nephropathy, and drug-induced kidney injury,^3^ in addition to mitochondrial disorders.^2^ Therefore, insights into kidney energy metabolism and mitochondrial function are crucial for understanding the mechanism of renal disease.

The kidney nephron is composed of various segments and cells. Previous studies have shown the possible contribution of different ATP synthesis pathways in each nephron segment^6^ based on the analysis of metabolic enzyme expression,^7,8^ electron microscopic findings,^9,10^ and metabolic analysis.^10^ Recently, we established GO-ATeam2 mice, which enabled the visualization of intracellular ATP dynamics at a single-cell level.^11^ Using this mouse line and two-photon microscopy, we revealed the crucial difference in intracellular ATP dynamics between proximal tubules (PTs) and distal tubules (DTs) during ischemia reperfusion (IR) injury,^11^ indicating the importance of direct visualization of the energy dynamics of individual nephron segments.

However, *in vivo* ATP imaging techniques have some critical limitations. First, deeper kidney regions, such as S3 segments of PTs and the medulla, cannot be observed from the kidney surface because the imaging depth of two-photon microscopy in the kidney is only 150 µm. Second, ATP synthesis inhibitors cannot be used in *in vivo* experiments because of their significant effects on circulation.

In the present study, we established a novel *ex vivo* observation system using a slice culture of GO-ATeam2 mouse kidneys, which enabled the visualization of intracellular ATP dynamics in various renal cells, including deeper nephron segments. We confirmed the viability of the kidney slices and subsequently revealed the ATP synthesis pathway of each nephron segment. We also visualized intracellular ATP dynamics in an *ex vivo* IR model and a cisplatin nephropathy model, and tested the efficacy of the candidate drug for mitochondrial protection.

Late last year, the FDA has eliminated animal testing requirements for drug approvals.^12^ This allows for the substitution of animal testing with experiments such organ-on-chip and organoids. The rationale behind this is animal protection and the inadequacy of animal models. However, at present, alternative methods have not yet been fully developed and are still awaiting further development. Our system retains a three-dimensional structure and a high degree of *in vivo* cellular functionality (e.g., drug uptake by transporters) and can also respond to disease models as *in vivo*. Additionally, by obtaining several slices from a single kidney, multiple experimental conditions can be tested simultaneously, thereby reducing the number of animals used, which is desirable from the perspective of animal welfare.

## Methods

### Preparation of kidney slices

After adequate anesthesia with isoflurane, the kidneys of 10–13-week-old male GO-ATeam2 mice were harvested and decapsulated. They were immediately sliced at 300 µm using a tissue slicer (VT1000S; Leica Microsystems, Wetzlar, Germany) in an ice-cold buffer (97.5 mM NaCl, 5 mM KCl, 0.24 mM NaH_2_PO_4,_ 0.96 mM Na_2_HPO_4_, 10 mM CH_3_COONa, 25 mM NaHCO_3_, 10 mM glucose, 5 mM Na pyruvate, 1.2 mM MgSO_4,_ and 1 mM CaCl_2_) gassed with 95% O_2_ and 5% CO_2_, as previously reported with a few modifications.^13^ As previously described, the slices were incubated at room temperature for 30 min before the experiments to detach dead or damaged cells loosely attached to the slice surface.^14^

### Ex vivo imaging setting

Kidney slices were placed in our original chamber and secured using a slice anchor. The buffer was gassed with 95% O_2_ and 5% CO_2_ during observation. Our system included a lens heater and a plate heater, and the buffer temperature was maintained at 37°C during all experimental procedures.

### Ex vivo IR injury model

To induce ischemia-like conditions, we replaced the normal buffer with a buffer with no energy sources and saturated it with nitrogen, as previously reported.^15,16^ During reperfusion, this buffer was replaced with the original buffer and oxygenation was resumed (Figure 3a).

### Analysis of intracellular ATP dynamics

The maximum FRET ratio range in each segment was defined as follows: pre FRET ratio − basal FRET ratio after the simultaneous administration of oligomycin and 2DG. The % ATP depletion was defined as follows: (pre FRET ratio – FRET ratio in each observation point) × 100/maximum FRET ratio range.

Full methods including image processing, mouse treatment, renal histological analysis, renal immunostaining are available in the Supplementary Methods.

## Results

### Establishment of the novel ATP imaging system with kidney slice culture

A FRET-based fluorescent ATP probe, GO-ATeam2, was developed for ATP imaging at single-cell resolution.^17^ This biosensor changes its structure depending on the intracellular ATP concentration, and had the linear correlation between the FRET ratio (Kusabira OFP signal/GFP signal) and the ATP concentration from 0.1 mM to 6.0 mM, making it possible to quantitatively measure the intracellular ATP concentration within the assumed range of variation.(Figure 1a).^17–19^ This ATP probe is also largely insensitive to pH changes within the physiologic ranges.^19^

**Figure 1.**
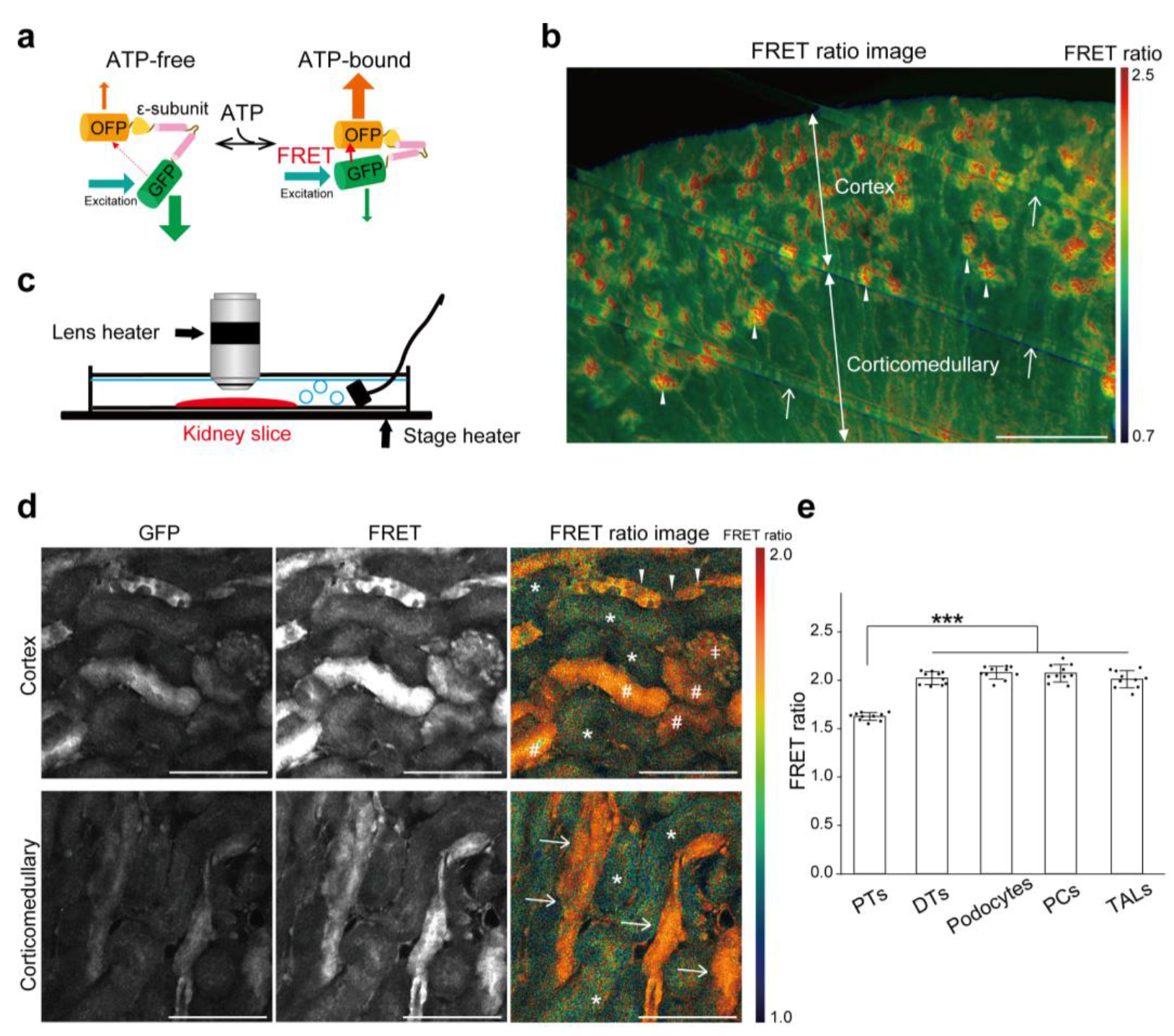
Establishment of the novel ATP imaging system using a kidney slice culture. (a) Schematic drawing of the ATP FRET biosensor. The ATP FRET biosensor changes its structure depending on the intracellular ATP concentrations. (b) ATP distribution shown by the ratio image of FRET/GFP emission in kidney slices using fluorescence stereomicroscopy. Arrowheads indicate glomeruli, and arrows indicate slice anchor strings. (c) Schematic drawing of the ATP imaging system using a kidney slice culture and two-photon microscopy. Briefly, 300-µm-thick kidney slices from GO-ATeam2 mice were placed into a chamber and secured with a slice anchor. (d) Visualization of intracellular ATP levels in PT (*), DTs (#), glomeruli (ǂ), CDs (arrowheads) and TALs (arrows). (e) FRET ratios (FRET/GFP) in PTs, DTs, podocytes, PCs and TALs (*n* = 10 slices). Statistical significance among nephron segments was assessed using one-way analysis of variance (ANOVA) with Tukey–Kramer *post hoc* tests for comparisons. *** *p* < 0.001. Scale bars: (b) 500 µm, (d) 100 µm.

Recently, we generated a novel mouse line systemically expressing GO-ATeam2 (GO-ATeam2 mice).^11^ We prepared kidney slices from GO-Ateam2 mice and established a novel *ex vivo* ATP imaging system that allowed us to observe all nephron segments simultaneously using fluorescence stereomicroscopy (Figure 1b). Detailed images of each segment could also be observed using two-photon microscopy (Figure 1, c and d). Warm colors indicate high FRET ratios (high ATP levels) and cool colors indicate low FRET ratios (low ATP levels) (Figure 1, b and d). Using nephron segment-specific antibodies, we identified PTs, DTs, thick ascending limbs (TALs) (Supplementary Figure S1a), and principal cells (PCs) of collecting ducts (CDs) (Supplementary Figure S1b). For PTs, PTs in the cortex (S1 and S2 segments) were analyzed, FRET, fluorescence resonance energy transfer; PTs, proximal tubules; DTs, distal tubules; CDs, collecting ducts; TALs, thick ascending limbs; PCs, principal cells.

except for the experiment in Supplementary Figure S5 where S3 segment was analyzed. Among glomerular cells, podocytes showed the highest biosensor expression (Supplementary Figure S1c). We assume that the differences in the expression of the biosensor among cell types are due to the different activities of the CAG promoter among cell types, which is knocked in the *R26* locus to enhance FRET probe expressison.^11^ The excitation laser was adjusted for each segment as previously described (PTs; high laser. DTs, TALs, PCs, and glomeruli [podocytes]; low laser).^11^ The FRET ratios in each segment (Figure 1e) were similar to those *in vivo*,^11^ suggesting that energy metabolism similar to that *in vivo* might be maintained in this slice culture system. Notably, cell-to-cell variability in intracellular ATP levels was relatively small in each segment (Figure 1e). To confirm the viability of the kidney cells, we performed the ATP imaging and the morphological evaluation at longer time points after the kidney slice preparation. At 24 h after the slice preparation, we observed the ATP decline and histological changes in some nephron segments (Supplementary Figure S2, a and b), but no apparent changes were detected at least at 6 and 12 h. In this study, all experiments were performed within 6 h after the kidney slice preparation.

***Intracellular ATP levels in each nephron segment after ATP synthesis inhibitor administration*** Elucidating the ATP synthesis pathways in each segment is important for understanding intracellular ATP dynamics under pathophysiological conditions. To inhibit ATP synthesis, we applied oligomycin A (oligomycin), an OXPHOS inhibitor, 2DG, a glycolytic inhibitor, and phloretin, a glucose transporters (GLUTs) inhibitor, to kidney slices. Although there was no apparent intracellular ATP change in all segments in vehicle-treated controls for 60 min (Supplementary Figure S3), simultaneous administration of oligomycin and 2DG decreased ATP levels in all segments to the basal levels within approximately 10 min (Figure 2, a–d), indicating that the administration of these drugs could completely inhibit ATP synthesis pathways.

**Figure 2.**
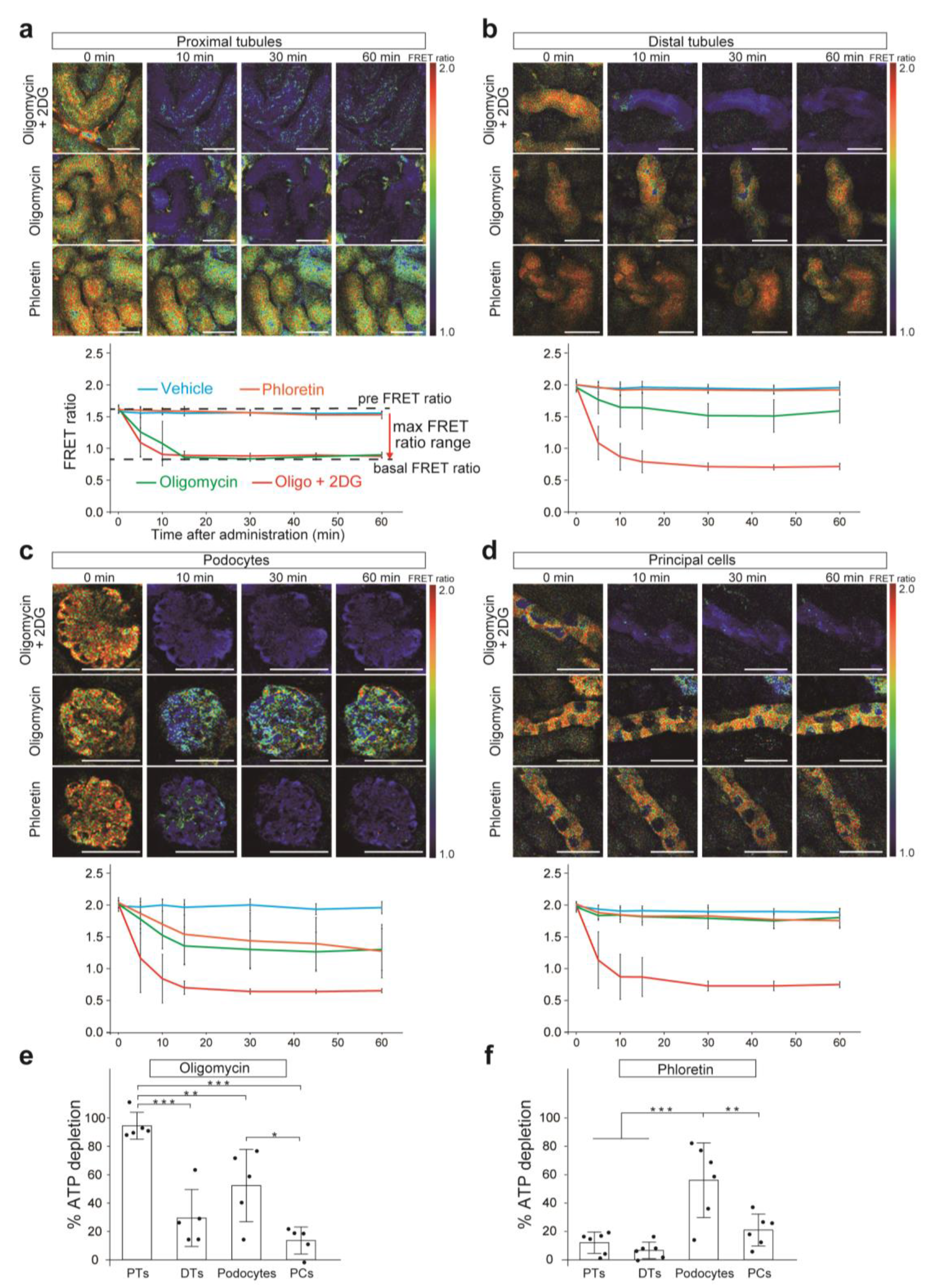
Intracellular ATP levels in each nephron segment after the administration of ATP synthesis inhibitors. (a–d) FRET ratio images in each nephron segment after the administration of oligomycin and 2DG (upper rows), oligomycin alone (middle rows), and phloretin alone (lower rows). Trends of FRET ratios are shown in graphs for each nephron segment after the administration of oligomycin and 2DG (red, *n* = 5 slices), oligomycin alone (green, *n* = 5 slices), phloretin alone (orange, *n* = 6 slices), and vehicle control (blue, *n* = 4 slices). ATP level in all segments reached the basal level after the administration of oligomycin and 2DG. The maximum FRET ratio range in each segment is defined as follows: (pre FRET ratio) − (basal FRET ratio after the administration of oligomycin and 2DG). (e, f) The % ATP depletion after the administration of oligomycin alone and phloretin alone in each nephron segment was shown. The % ATP depletion was defined as follows: [(pre FRET ratio) − (FRET ratio after the administration of each inhibitor for 60 min)] × 100 (%)/(maximum FRET ratio range). Statistical significance among nephron segments was assessed using one-way analysis of variance (ANOVA) with Tukey–Kramer *post hoc* tests for comparisons. * *p* < 0.05; ** *p* < 0.01; *** *p* < 0.001. Scale bars: 50 µm. 2DG, 2-deoxy-D-glucose; PTs, proximal tubules; DTs, distal tubules; PCs, principal cells.

**Figure 3.**
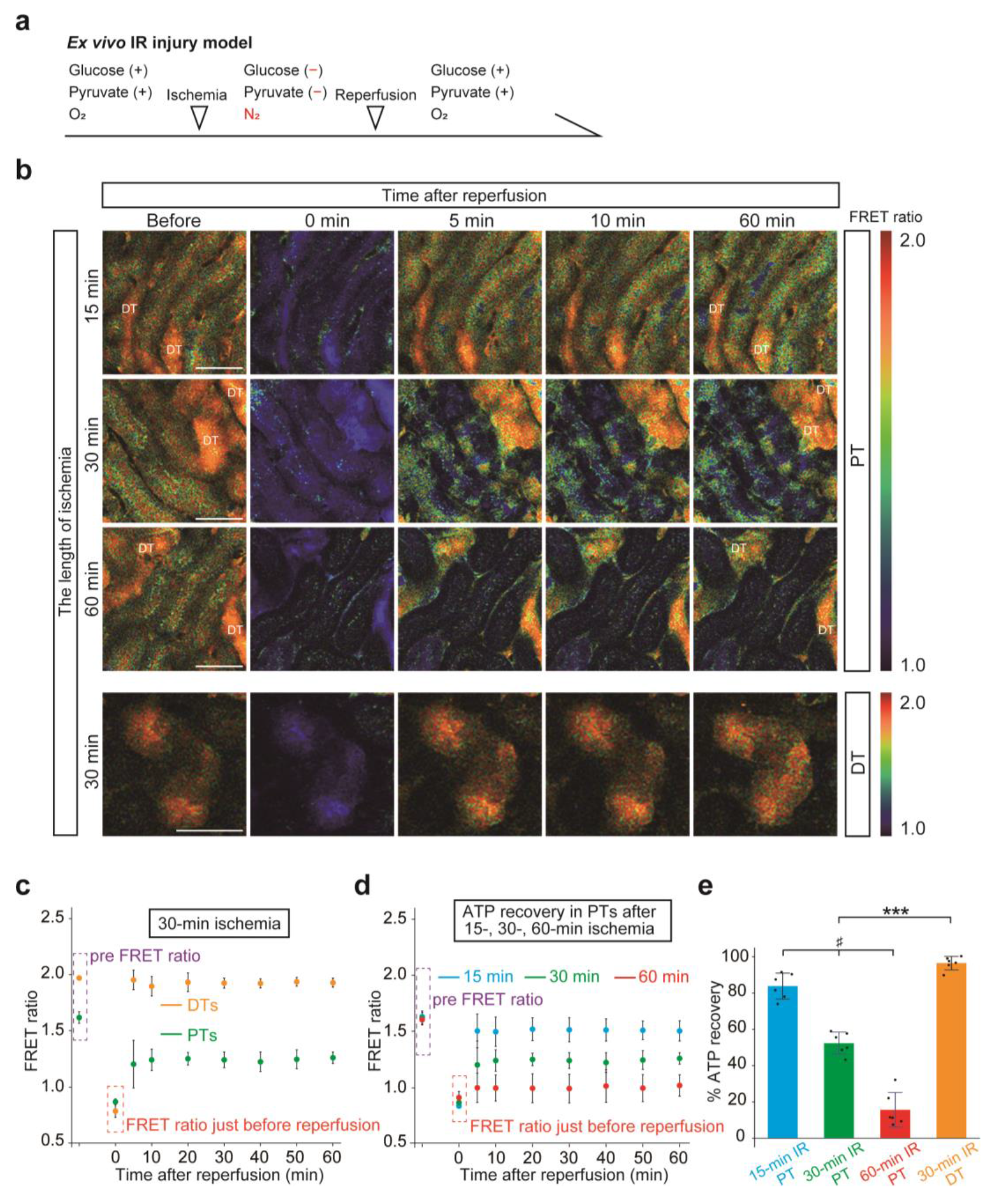
ATP recovery in PTs varies depending on the length of ischemia in an *ex vivo* IR injury model. (a) Schematic drawing of *ex vivo* IR injury model. (b) FRET ratio images of PTs during reperfusion after 15-, 30-, and 60-min ischemia and those of DTs after 30-min ischemia. (c) FRET ratio graph of PTs (green) and DTs (yellow) during reperfusion after 30-min ischemia (*n* = 6 slices). (d) FRET ratio graph of PTs during reperfusion after 15-, 30-, and 60-min ischemia (blue, green, and red, respectively). (*n* = 6 kidney slices per groups). FRET ratio graph of PTs after 30-min ischemia in (c) are presented here again. (e) The % ATP recovery is defined as follows: [(FRET ratio 60 min after reperfusion) − (FRET ratio just before reperfusion)] × 100 (%)/{(pre FRET ratio) − (FRET ratio just before reperfusion)}. Statistical significance was assessed using trend test across 15-, 30-, and 60-min IR groups. Differences between the two groups (30-min IR PT versus 30-min IR DT) were compared using unpaired, two-tailed *t*-test. # *p* for trend < 0.001; *** *p* < 0.001. Scale bars: 50 µm. IR, ischemia–reperfusion; PTs, proximal tubules; DTs, distal tubules.

We then analyzed intracellular ATP dynamics after administering oligomycin alone and found intriguing differences in the ATP decline rate and ATP plateau levels among nephron segments. In PTs, the ATP decline rate and plateau levels were similar to those observed after simultaneous administration of oligomycin and 2DG (Figure 2a). The % ATP depletion, which was defined as (pre FRET ratio – FRET ratio in each observation point) × 100/ (pre FRET ratio − basal FRET ratio after simultaneous administration of oligomycin and 2DG) as detailed in the Methods section, was 94.5% at 60 min (Figure 2e), indicating a high dependence of PTs on OXPHOS. Conversely, ATP levels in DTs and PCs decreased slowly and moderately (Figure 2, b and d), and the % ATP depletions were only 29.4% and 13.5%, respectively (Figure 2e). These findings are consistent with those of our previous *in vivo* study, which showed that ATP levels in DTs decreased more slowly than those in PTs after ischemia. In podocytes, oligomycin administration reduced ATP levels to 47.7% (Figure 2, c and e), suggesting that podocytes require moderate OXPHOS for ATP synthesis.

To examine the effect of glycolysis on ATP synthesis, we then administered 2DG alone, but did not find any marked change in any segment. Even at a concentration of 100 mM, only the ATP levels in podocytes decreased slightly (Supplementary Figure S4, a–d). Because rapid and significant ATP depletion was observed in all segments with the simultaneous administration of oligomycin and 2DG, we could not elucidate the precise reason for the ineffectiveness of 2DG alone. Instead, we administered phloretin, a GLUTs inhibitor, and found that podocytes, but not other segments, showed a significant reduction in ATP levels (Figure 2c), and the % ATP depletion in podocytes was 56.1% (Figure 2f). This finding indicates that podocytes actively take up glucose via GLUTs and utilize both OXPHOS and glycolysis for ATP synthesis.

***IR injury in an ex vivo slice culture system recapitulates intracellular ATP dynamics for in vivo IR injury*** To confirm whether this novel *ex vivo* system could recapitulate intracellular ATP dynamics in kidney diseases, we first induced culture conditions that mimicked IR injury^15,16^ and assessed intracellular ATP dynamics (Figure 3a; see the Methods section for details). The ATP levels in both PTs and DTs decreased rapidly to the basal levels after 30-min ischemia, but recovered differently after reperfusion (Figure 3, b and c). The % ATP recoveries in PTs and DTs 60 min after reperfusion were 52.3% and 97.9%, respectively (Figure 3, b, c and e), indicating the resistance of DTs to IR injury. Next, we analyzed intracellular ATP dynamics in PTs after different ischemic times. A longer ischemic time resulted in poorer ATP recovery in PTs (Figure 3, b and d). The % ATP recovery in PTs 60 minutes after reperfusion was 83.8% after 15- minute ischemia, 52.3% after 30-minute ischemia, and 15.6% after 60-minute ischemia (Figure 3e).

Electron microscopy analysis of these slices as early as 1 h after reperfusion revealed mitochondrial damage in PTs (Figure 4a) and a lower mitochondrial length-to-width ratio (L/W ratio) after longer ischemia (Figure 4b). Additionally, the slices 6 h after reperfusion showed loss of brush borders in PTs and tubular degeneration after longer ischemia (Figure 4, c and d). Taken together, this slice culture system could recapitulate intracellular ATP dynamics during IR injury *in vivo*, as ATP recovery in DTs was better than that in PTs after reperfusion, and PT recovery was worse after prolonged ischemia.^11^ This system also faithfully recapitulated mitochondrial damage and histological changes after prolonged ischemia, suggesting its reliability. We also evaluated intracellular ATP dynamics of PTs in the corticomedullary region during IR injury, which could not be observed in our previous *in vivo* observation from the kidney surface. Whereas the FRET ratio in PTs in the corticomedullary region was slightly lower than that in PTs in the cortex (Supplementary Figure S5b), intracellular ATP dynamics and the % ATP recovery were comparable between groups (Supplementary Figure S5, a, c and d).

**Figure 4.**
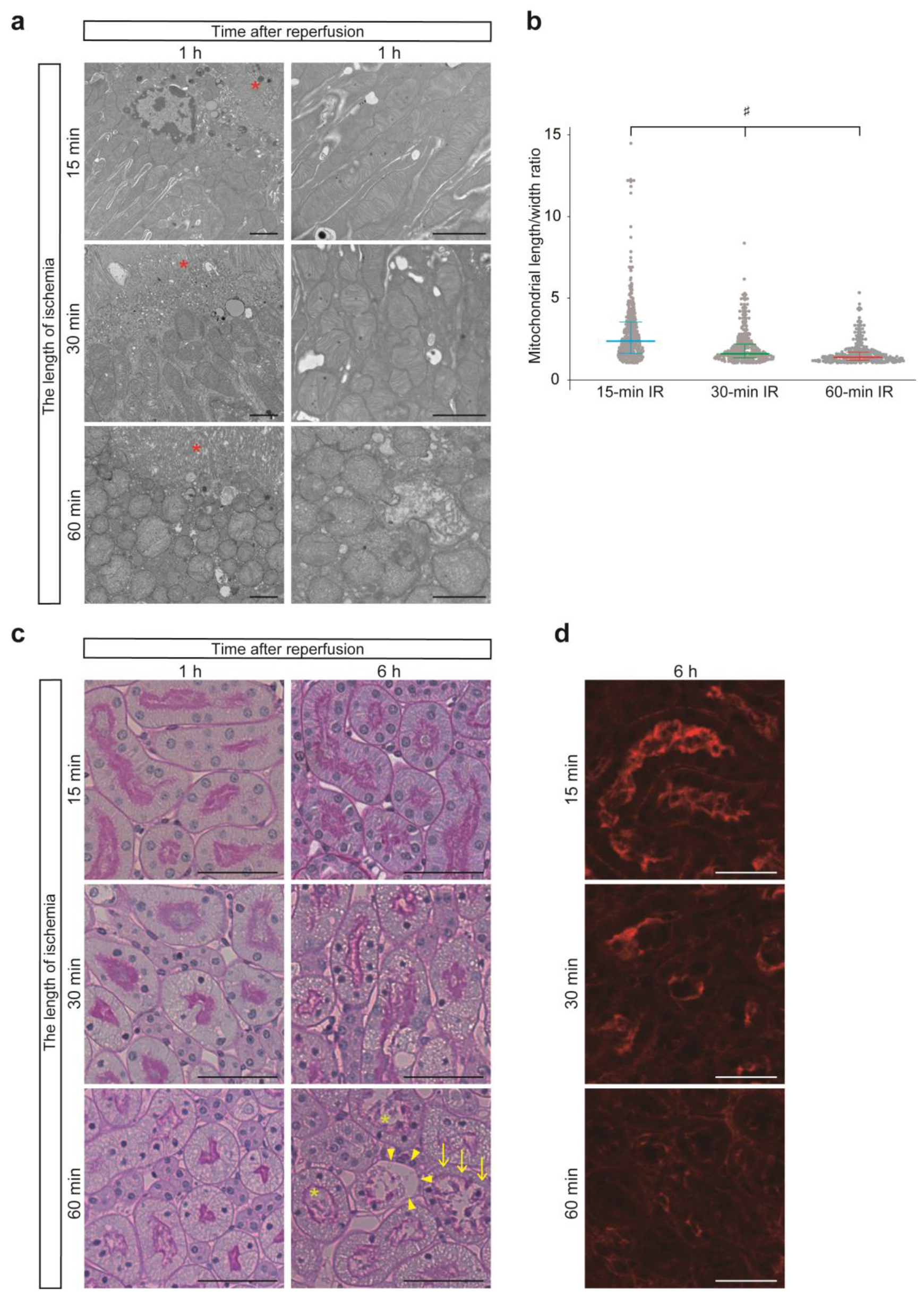
Histological changes in *ex vivo* IR injury model. (a) At 1h after reperfusion, electron microscopy analysis showed that a longer ischemic time resulted in more pronounced changes in the mitochondrial structures in PTs. *: brush borders. (b) Quantitative evaluation was performed using the mitochondrial length/width ratio (*n* = 3 slices, 9 tubules, 450 mitochondria). (c) PAS staining of the kidney slices 1 h and 6 h after reperfusion. At 1 h after reperfusion, no apparent histological injury or only a minor change was observed, even in the slices that underwent 60-min ischemia. At 6 h after reperfusion, tubular injuries, such as tubular epithelial shedding (arrowheads), debris (*), and brush border loss (arrows) were observed in the slices that underwent a longer ischemia. Representative images are shown. (d) Immunostaining of the slices 6 h after 15-, 30-, 60-min IR. Positive phalloidin staining indicates brush borders in PTs. A healthy tall brush border was maintained in kidney slices underwent 15-min IR, whereas a shedding or shortened brush border was observed in the slices underwent 60-min IR. Representative images are shown. Statistical significance was assessed using trend test across 15-, 30-, and 60-min IR groups (b). # *p* for trend < 0.001. Scale bars: (a) 2 µm, (c) 50 µm, (d) 25 µm. IR, ischemia reperfusion; PTs, proximal tubules; PAS, Periodic acid–Schiff.

### Mitochondrial dysfunction and ATP depletion in cisplatin nephropathy in vivo

Next, we analyzed intracellular ATP dynamics in a cisplatin nephropathy model in which mitochondrial dysfunction plays an important role.^20^ We first evaluated intracellular ATP dynamics in cisplatin nephropathy *in vivo* and found ATP-depleted cells in degenerated tubules (Figure 5a), although the segments could not be identified due to severe injury. Mitochondrial damage in cisplatin nephropathy was further confirmed by lower mitochondrial L/W ratios (Figure 5b), and reduced mitochondrial membrane potential was evaluated using TMRM analysis (Figure 5c). From these findings, we confirmed tubular ATP depletion in the cisplatin nephropathy model, possibly due to mitochondrial damage. However, it was still unclear which nephron segments were damaged and whether there was a way to alleviate damage.

**Figure 5.**
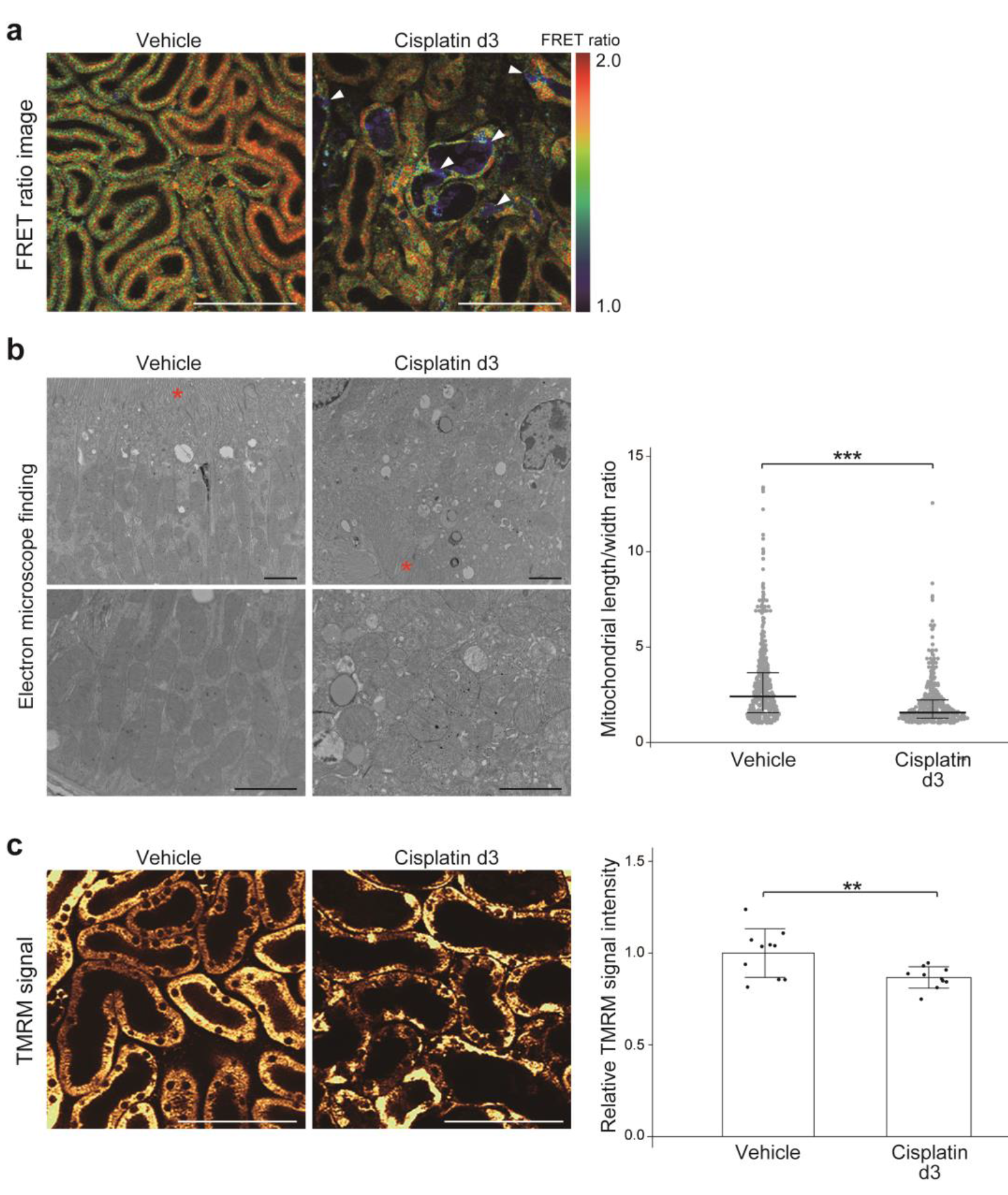
**Mitochondrial dysfunction and ATP depletion in cisplatin nephropathy *in vivo.*** (a) FRET ratio images in living kidneys of mice that were administered cisplatin. Mice were administered 15 mg/kg of cisplatin or vehicle intraperitoneally 3 days before the observation. In the cisplatin group, ATP-depleted cells (arrowheads) were detected in damaged tubules and debris was confirmed in the lumens. Identification of the segments was technically challenging because of the morphological changes. (b) Mitochondrial damage was detected in PTs of the cisplatin group. Quantitative evaluation was performed using the mitochondrial length/width ratio (*n* = 3 mice, 9 tubules, 450 mitochondria). *: brush borders. (c) Mitochondrial membrane potential shown by the administration of TMRM was reduced in the kidneys of cisplatin-treated mice (*n* = 3 mice, total of 10 views in each group). Statistical significance was assessed using unpaired, two-tailed *t*-test. ** *p* < 0.01; *** *p* < 0.001. Scale bars: (a, c) 100 µm, (b) 2µm. PTs, proximal tubules; TMRM, tetramethyl rhodamine methyl ester.

### ATP levels decrease both in PTs and DTs in a cisplatin nephropathy model

To answer this question, we evaluated the sensitivity of various nephron segments to cisplatin, utilizing this *ex vivo* ATP imaging system. Several concentrations of cisplatin were administered referring to the previous report using kidney slices.^21^ After cisplatin administration, the ATP levels decreased mainly in PTs and DTs, but not in podocytes or PCs (Figure 6a). The administration of a higher concentration of cisplatin resulted in more rapid and severe ATP depletion in PTs and DTs (Figure 6, b and c).

**Figure 6.**
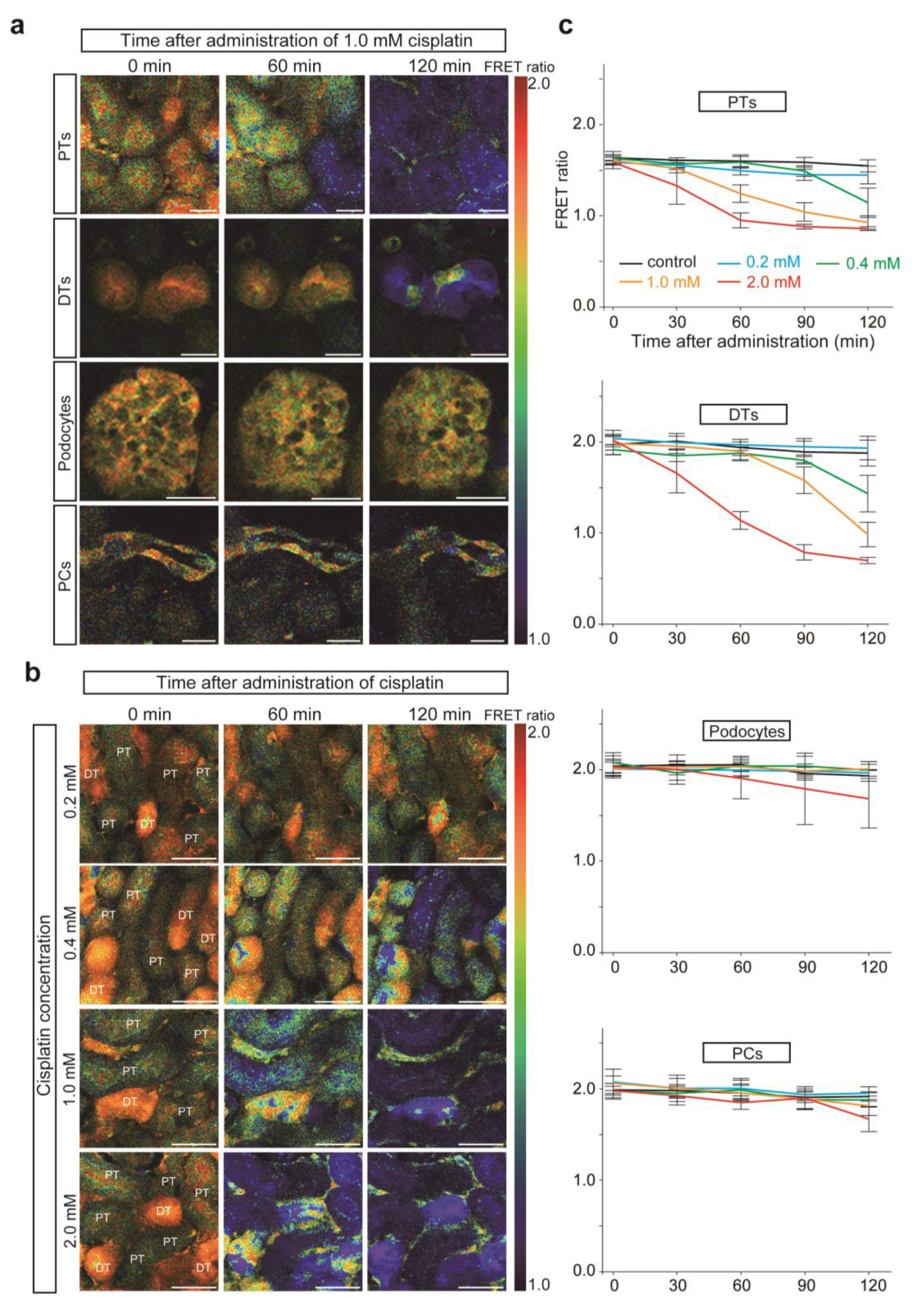
Sensitivity to cisplatin differs among the nephron segments. (a) FRET ratio images after the administration of 1.0 mM cisplatin. During 120 min of observation, ATP levels in PTs and DTs, but not those in podocytes and PCs, decreased significantly. (b) FRET ratio images after the administration of different concentrations of cisplatin. (c) FRET ratio graphs after the administration of different concentrations of cisplatin (*n* = 5 slices in 0.2 mM, 0.4 mM, 1.0 mM groups and *n* = 6 slices in control and 2.0 mM groups). Scale bars: (a) 25 µm, (b) 50 µm. PTs, proximal tubules; DTs, distal tubules; PCs, principal cells.

### Cisplatin nephrotoxicity is mediated via OCT2 in PTs but not in DTs in an ex vivo system

Cisplatin accumulates within tubules via cellular uptake by OCT2, which is specifically expressed on the basolateral side of PTs,^22,23^ explaining its nephrotoxicity in PTs. In our analysis utilizing an *ex vivo* ATP imaging system, however, ATP levels in DTs were also reduced. To test whether cisplatin also causes DT damage, we treated wild-type mice with cisplatin. In addition to the upregulation of *Kim1* mRNA, we found that the expression of kidney *Neutrophil gelatinase-associated lipocalin* (*Ngal*) mRNA was increased (Supplementary Figure S6a), and the NGAL protein appeared in DTs (Supplementary Figure S6b), suggesting DT damage.^24^ NGAL staining in PTs was considered as reabsorbed NGAL from the urine. Therefore, we examined the mechanisms of ATP depletion induced by cisplatin in PTs and DTs using this *ex vivo* system. First, the expression levels of OCT2 protein and mRNA in kidney slices in this *ex vivo* system were comparable with those of whole kidneys (Figure 7, a and b). Next, we confirmed using the ICP-MS method that cisplatin was taken up by the slices, which was attenuated by the simultaneous administration of cimetidine, an OCT2 inhibitor^25,26^ (Figure 7c). Electron microscopy findings also showed that the mitochondrial L/W ratio of PTs was reduced by the administration of cisplatin, which was reversed by the administration of cimetidine (Figure 7, d and e). These results indicate OCT2-mediated cisplatin uptake and its inhibition by cimetidine in this *ex vivo* system.

**Figure 7.**
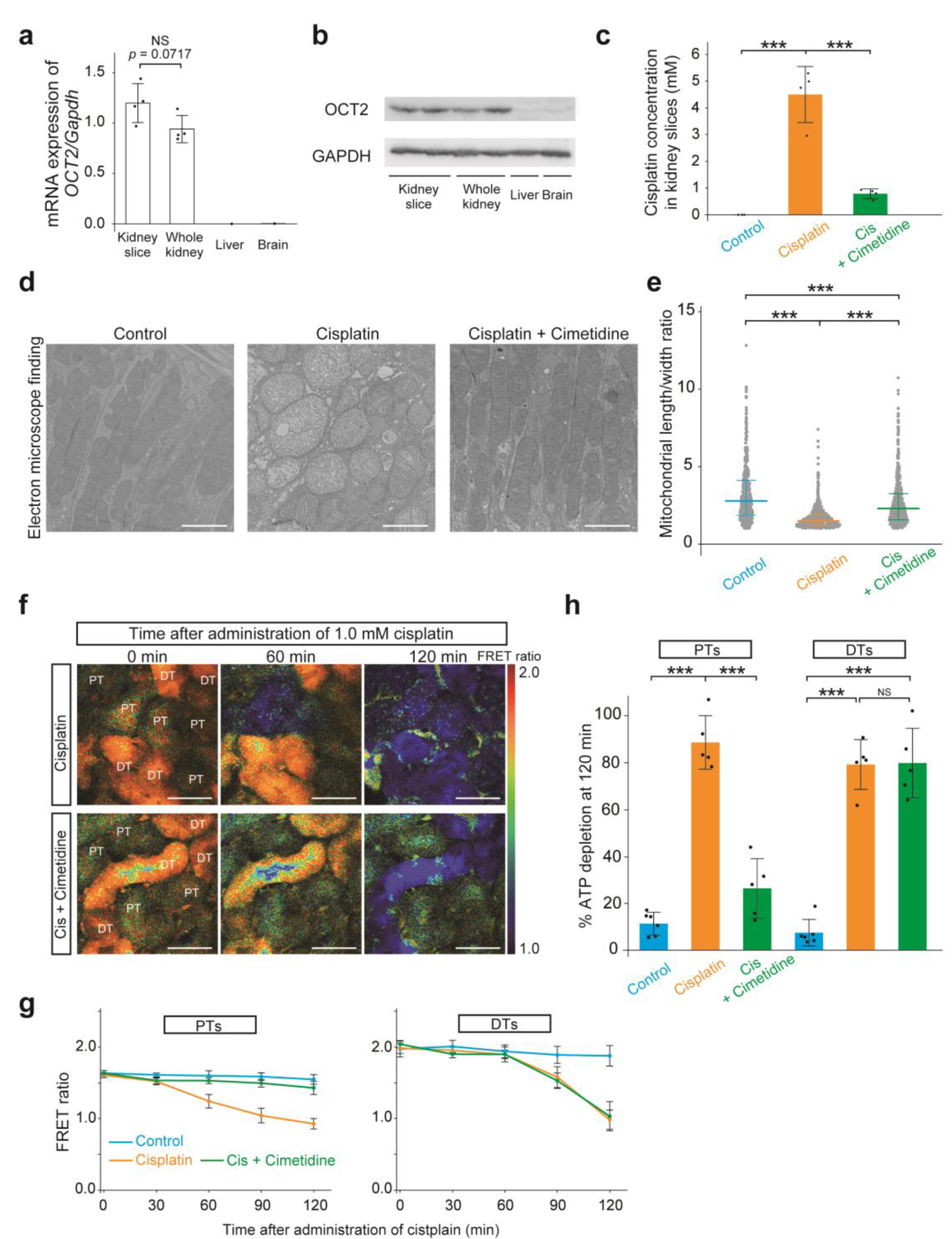
Cisplatin nephrotoxicity in PTs, but not that in DTs, is OCT2 mediated. (a, b) *OCT2* mRNA and OCT2 protein expression levels in kidney slices. Both levels were similar between kidney slices and whole kidneys (*n* = 4 per group in Figure 7a). Mouse liver and brain samples were used as negative controls. (c) The amount of cisplatin uptake in kidney slices was determined using ICP-MS method with or without cimetidine (an OCT2 inhibitor) (*n* = 3 slices in the control group and *n* = 4 slices in the cisplatin [1.0 mM] and the cisplatin [1.0 mM] + cimetidine [1.0 mM] groups). (d) Mitochondrial morphological changes were obvious in PTs in the cisplatin group, which was attenuated in PTs in the cisplatin + cimetidine group. (e) Quantitative evaluation of mitochondria morphology using the mitochondrial length/width ratio (*n* = 3 slices, 9 tubules, 450 mitochondria per each group). (f) FRET ratio images after the administration of cisplatin and cimetidine. (g, h) FRET ratio graphs in PTs and DTs during 120 min of observation (g), and the % ATP depletion in PTs and DTs 120 min after the administration of reagents (h). FRET ratio graphs in the control and 1.0 mM cisplatin groups in Figure 6c are presented again for comparison (*n* = 5 slices per group). Statistical significance was assessed using one-way analysis of variance (ANOVA) with Tukey–Kramer *post hoc* tests (c, e, and h) or unpaired, two-tailed *t*-test (a) for comparisons. *** *p* < 0.001; NS, not significant. Scale bars: (d) 2 µm, (f) 50 µm. PTs, proximal tubules; DTs, distal tubules; OCT2, organic cation transporter 2; ICP-MS, inductively coupled plasma-mass spectrometry.

Furthermore, cimetidine ameliorated cisplatin-induced ATP depletion in PTs but not in DTs (Figure 7, f– h). These results indicate that the mechanism of DT injury in cisplatin nephropathy may be OCT2 independent.

### Mitochondrial protective reagent ameliorates ATP depletion induced by cisplatin

Finally, we examined whether this *ex vivo* system is also useful for testing candidate drugs for mitochondrial protection. To validate this system, we evaluated intracellular ATP dynamics after cisplatin administration and simultaneous administration of MA-5, a mitochondria protection reagent^27^. MA-5 binds to mitofilin, which maintains mitochondrial cristae structure, increases ATP synthesis efficiency, and is also reported to ameliorate cisplatin-induced renal tubular damage.^28,29^ MA-5 treatment ameliorated ATP depletion induced by cisplatin compared with the vehicle group for as long as 120 min (Figure 8, a–c).

**Figure 8.**
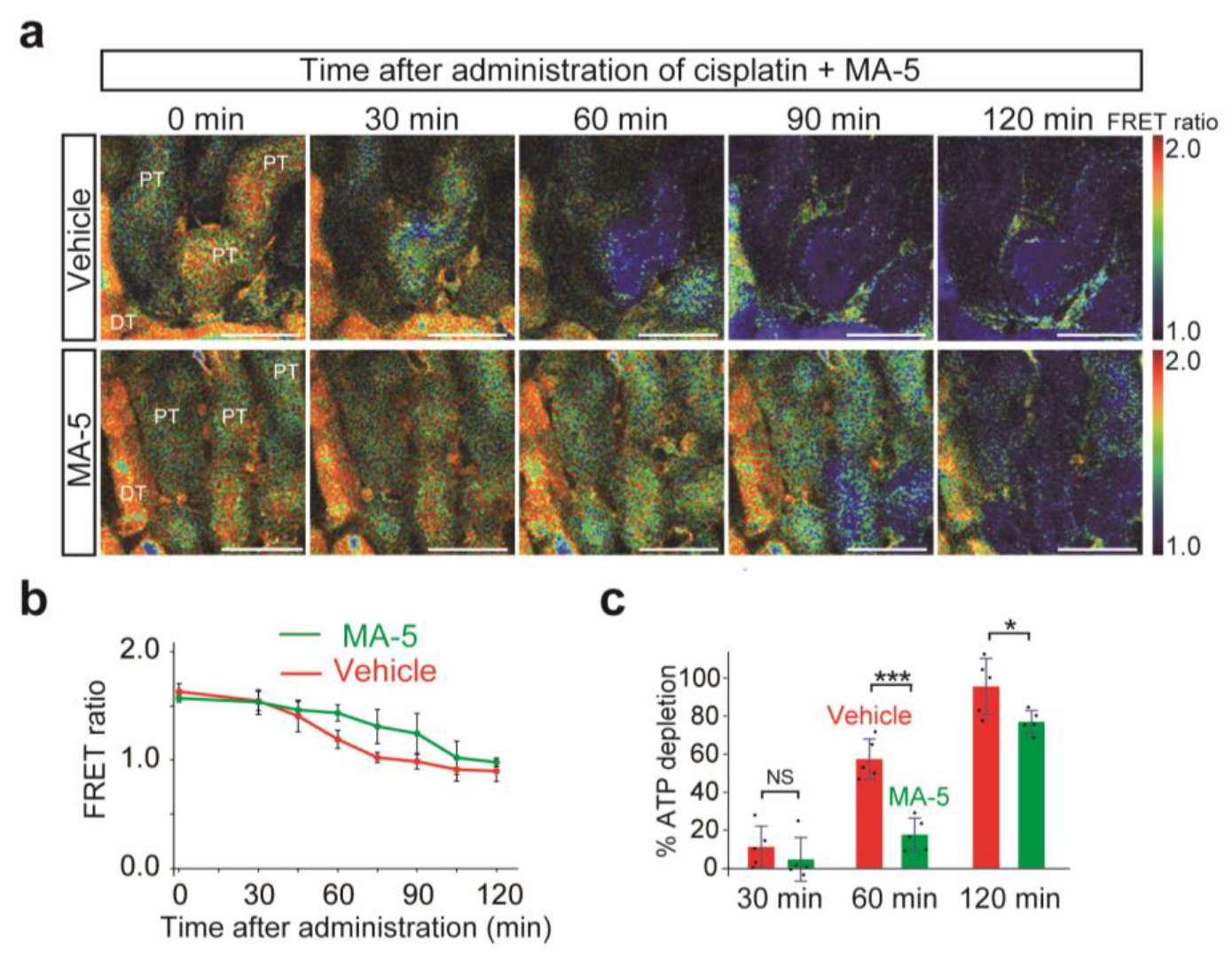
MA-5 treatment ameliorates ATP depletion induced by cisplatin. (a) FRET ratio images after the administration of cisplatin and MA-5 or vehicle. (b, c) FRET ratio graphs in PTs during 120 min of observation (b) and the % ATP depletion in PTs 30, 60 and 120 min after the administration of reagents (c) (*n* = 5 slices per group). Statistical significance was assessed using unpaired, two-tailed *t*-test. * *p* < 0.05; *** *p* < 0.001; NS, not significant. Scale bars: 50 µm. MA-5, mitochonic acid 5; PTs, proximal tubules; DTs, distal tubules.

## Discussion

We successfully established a novel ATP imaging system utilizing kidney slice culture, which enabled the evaluation of intracellular ATP dynamics in the whole kidney, including deeper nephron segments. Additionally, we have directly demonstrated that each nephron segment had different ATP synthesis pathways. We also applied an IR injury model, which could reproduce intracellular ATP dynamics *in vivo*, and a cisplatin nephropathy model, which showed different injury mechanisms between PTs and DTs, in this *ex vivo* system. Finally, we tested the efficiency of a mitochondrial protection reagent, which delayed cisplatin-induced ATP decrease.

The *ex vivo* slice culture system showed preserved three-dimensional structure, cell–cell interactions, and functional transporters, including OCT2.^14,30,31^ This culture system better reflects the pathophysiological conditions compared with *in vitro* cell culture studies or *ex vivo* studies using micro-dissected nephron segments. ATP visualization of whole kidneys using this *ex vivo* system is a powerful tool for understanding energy dynamics in the kidneys.

In our previous study, we demonstrated the crucial difference in intracellular ATP dynamics between PTs and DTs, and found that PTs were more susceptible to ischemia than DTs.^11^ However, the mechanism of ATP synthesis remained unclear. Our novel *ex vivo* ATP imaging system enabled us to assess these cells accurately by administering OXPHOS and glycolytic inhibitors. After the administration of oligomycin alone, ATP levels in PTs rapidly decreased to the basal levels, indicating that the glycolytic pathway did not significantly contribute to ATP synthesis in PTs (Figure 2, a, e and f). Previous reports have shown that glycolytic enzyme expression in PT is very low,^7,8^ which is consistent with our results. Furthermore, our data showed directly that the glycolytic pathway was not activated, even when OXPHOS was completely inhibited.

Podocyte energy metabolism has been controversial. Whereas mitochondrial density in podocytes is low, and anaerobic glycolysis has been predominant in podocyte metabolic pathways,^10^ another group reported significant involvement of the OXPHOS pathway in podocyte energy homeostasis.^32^ In our present study, we demonstrated that, in podocytes, GLUTs-mediated glucose uptake was considerably active, which was followed by the glycolysis pathway for ATP synthesis, whereas the OXPHOS pathway was also important (Figure 2, c, e and f). Conversely, ATP levels in DTs and PCs were maintained for 60 min after administration of oligomycin or phloretin alone, indicating that both the OXPHOS and glycolysis pathways could be activated in these segments when one pathway was inhibited (Figure 2, b, d–f). This mechanism might be why these segments are more resistant to IR injury.^33^

We next applied IR injury to this system and, similar to our previous *in vivo* data,^11^ found that a longer ischemia worsened ATP recovery in PTs after reperfusion, whereas intracellular ATP dynamics in DTs were more resistant to an ischemic insult (Figure 3). Intracellular ATP depletion in PTs is known to lead to rapid disruption of the apical actin cytoskeleton structure and subsequent brush border loss *in vivo*.^34^ We also found shortened brush borders in this system after longer ischemia, in which ATP recovery was poor (Figure 4d). These results indicate that this novel *ex vivo* system can accurately recapitulate intracellular ATP dynamics and pathological changes *in vivo*. In a mouse IR model, PTs in the corticomedullary region have been reported to be the most susceptible.^35–37^ We hypothesized that energy metabolism and ATP recovery in PTs in the corticomedullary region could be different from those in the cortex. However, in contrast to our expectations, we found that ATP recovery after reperfusion in PTs in both regions was comparable (Supplementary Figure S5). This similarity might be due to a lack of impaired blood flow after ischemia in this *ex vivo* system. Persistent impaired blood flow has been reported after IR injury, especially in the corticomedullary region.^38–41^ This finding might provide important insights into the reported differences in vulnerability between PTs in the corticomedullary region and the cortex, which could be due, at least partly, to the differences in hemodynamic alterations rather than in segment-specific metabolic properties.

Furthermore, we, for the first time, elucidated intracellular ATP dynamics in cisplatin-induced renal injury *in vivo* and in this *ex vivo* system (Figure 5, Figure 6 and Figure 7). We clearly demonstrated the ATP decline in DTs and PTs, which are the main site of injury in cisplatin nephropathy. There have been a few reports indicating that cisplatin could also damage DTs and cause hypomagnesemia in patients on cisplatin therapy.^42^ Increased expression of *Ngal* mRNA in the kidneys and localization of NGAL protein in distal tubules *in vivo* also supported DT injury by cisplatin (Supplementary Figure S6). We also demonstrated that cisplatin was taken up in the slices, and the administration of cimetidine, an OCT2 inhibitor, attenuated cisplatin uptake in the slices and ATP decline in PTs, but not in DTs (Figure 7, c–h). It is unclear whether this is because of a different mechanism of injury in DTs, or whether ATP decline in DTs is a secondary response to PT injury. However, the fact that cimetidine did not attenuate ATP decline in DTs, but almost entirely canceled ATP decline in PTs, makes it plausible to assume that ATP decline in DTs is not a secondary impairment after PT injury. Cisplatin transporters other than OCT2, such as CTR1,^43^ OAT1, and OAT3,^44^ have been postulated, and further study is warranted to determine whether other transporters are involved in cisplatin uptake in DTs. Cimetidine is also a histamine receptor 2 antagonist, and according to the KIT databases,^45^ histamine receptor 2 is expressed in PTs. Because renoprotective effects of histamine receptor 2 antagonists have reported,^46,47^ it is theoretically possible that preservation of ATP levels in PTs xafter cimetidine administration could be attributed at least in part due to its renoprotection. However, histamine release from circulating cells is considered negligible in our *ex vivo* system.

Our present study has several technical limitations as follows: (1) Although kidney slice preparation was performed promptly in an ice-cold buffer with oxygenation, it is challenging to avoid cellular damage completely. Apparently damaged kidney slices were excluded from the analysis in this study. (2) Although the kidney, especially the medullary region, is hypoxic *in vivo*,^48,49^ the aeration is required in this system. When aeration was turned off, we observed a rapid decrease in ATP levels, especially in PTs over time (Supplementary Figure S7). However, even under aeration, the basal ATP levels in each segment and the ATP dynamics in the IR injury model were very similar to those *in vivo*. In this regard, we consider that this system recapitulates intracellular ATP dynamics *in vivo*. (3) This system lacks blood and urine flow, influencing disease progression *in vivo*.

Despite these limitations, analysis utilizing the novel kidney slice culture ATP imaging system provides valuable information about the segment-specific ATP synthesis pathways and intracellular ATP dynamics in disease models, leading to a new understanding of disease mechanisms. This system is easy to use for controlled intervention experiments. It allows multiple slices to be cultured from a single kidney and various experimental conditions to be tested in one mouse, which is appropriate from an animal welfare perspective. Kidney slices from mouse models of diabetes, aging, drug-induced nephropathy, and genetic disorders could also provide useful experimental models for studying metabolic changes under these conditions. While we used pyruvate and glucose as energy substrates in these experiments, the effects of certain energy substrates on the kidney energy dynamics can be studied in detail by changing the culture buffer composition. This novel ATP imaging system is a powerful tool to evaluate renal energy metabolism in various kidney diseases and provides us with a deeper understanding of the mechanisms underlying kidney disease.

## Data Statement

All data used in this study are available in this article.

## Acknowledgments

This research was supported by the Japan Agency for Medical Research and Development (AMED) under grant numbers AMED-CREST 22gm1210009 (MoY), 21gm5010002 (MoY), 22zf0127003h001 (MoY), 22ek0310020h0001 (MoY), and 22zf0127001 (TaA); KAKENHI Grant-in-Aid for Scientific Research B (20H03697 to MoY) and Grant-in-Aid for Young Scientists (21K16162 to ShinY) from the Ministry of Education, Culture, Sports, Science, and Technology of Japan. This work was also supported by JSPS KAKENHI Grant Number JP16H06280, Grant-in-Aid for Scientific Research on Innovative Areas – Platforms for Advanced Technologies and Research Resources “Advanced Bioimaging Support”. This work was also supported by the grants from the Uehara Memorial Foundation (MoY), the Takeda Science Foundation (MoY), the Sumitomo Foundation (MoY), the Mochida Memorial Foundation for Medical and Pharmaceutical Research (ShinY) and Suzuken Memorial Foundation (ShinY).

The authors are grateful to Professor Michiyuki Matsuda for valuable suggestion and guidance. This work was supported partly by the World Premier International Research Center Initiative (WPI), MEXT, Japan and the Kyoto University Live Imaging Center. Part of this work was included as an abstract at the Annual Meeting of the American Society of Nephrology.

## Author Contributions

Shigenori Yamamoto and M. Yanagita designed the experiments and wrote the manuscript; M. Yanagita supervised the project; Shinya Yamamoto, M. Takahashi, A. Mii, A. Okubo, S. Nakagawa, M. Yamamoto and Shigenori Yamamoto performed experiments; S. Fukuma, N. Toriu, M. Yamamoto and Shigenori Yamamoto analyzed the data. T. Abe and H. Imamura provided the resources, reviewed and edited the manuscript.

