## Supplementary Figure S1 for "Visualization of intracellular ATP dynamics in the whole kidney under pathophysiological conditions using the kidney slice culture system"

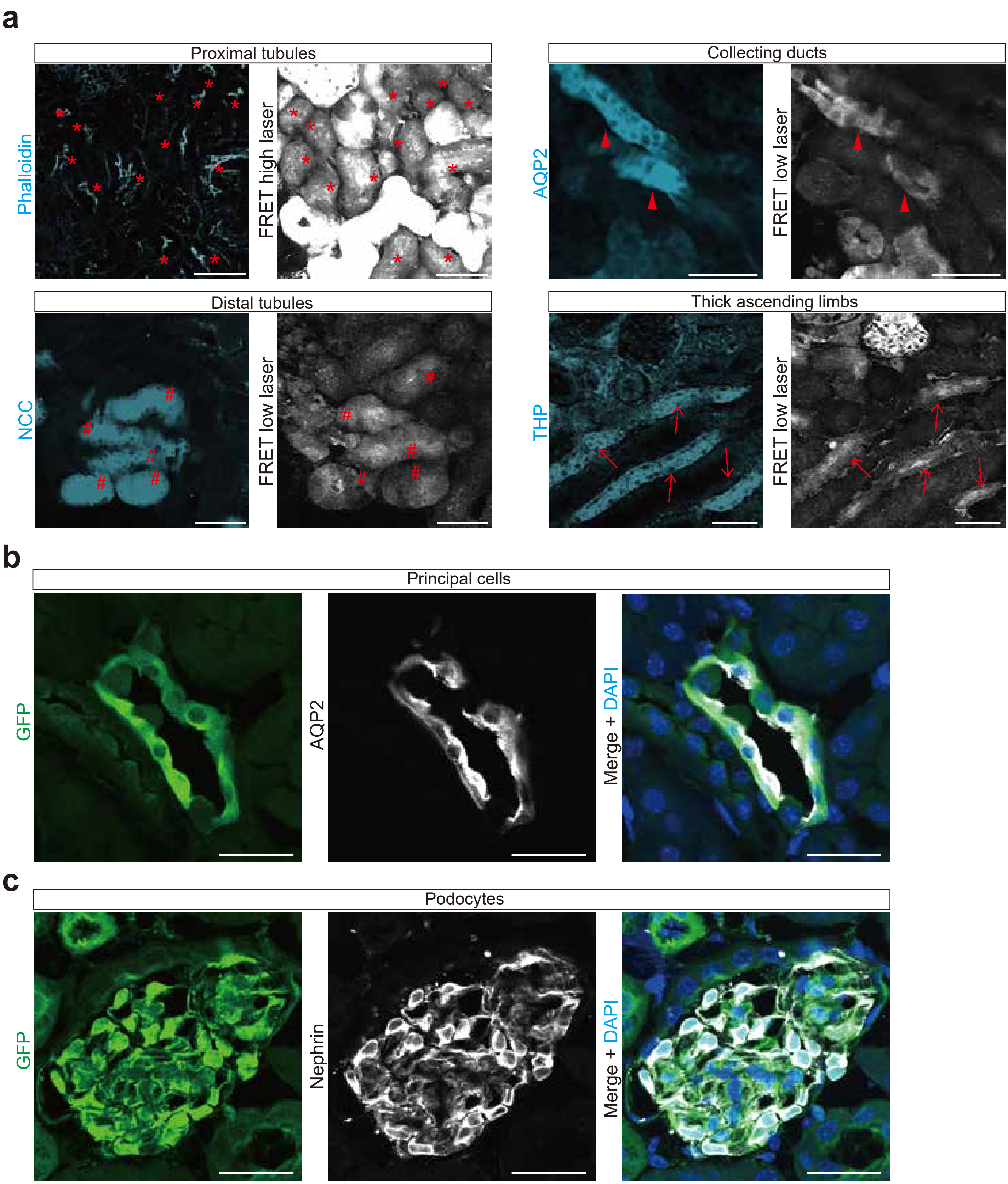


**Supplementary Figure S1. Identification of each nephron segment by immunostaining analysis.** (a) Identification of proximal tubules (*), distal tubules (#), thick ascending limbs (arrows), and collecting ducts (arrowheads) by immunostaining analysis. After the observation of FRET signals, kidney slices were fixed in 4% paraformaldehyde and directly immunostained with nephron segment-specific antibodies, so that tubules appear slightly smaller in the stained images than in the imaging. (b, c) Immunostaining of GFP, AQP2, and nephrin in the kidneys of GO-ATeam2 mice showed that cells with strong GO-ATeam2 expression in collecting ducts and glomeruli were mainly principal cells and podocytes, respectively. Other constituent cells, such as intercalated cells, mesangial cells, and endothelial cells, were not evaluated in this study due to weak GO-ATeam2 signals. Scale bars: (a) 50 μm. (b, c) 25 μm.

FRET, fluorescence resonance energy transfer; AQP2, aquaporin 2; NCC, sodium chloride co-transporter; THP, Tamm–Horsfall protein; GFP, green fluorescent protein.
