## Supplementary Figure S2 for "Visualization of intracellular ATP dynamics in the whole kidney under pathophysiological conditions using the kidney slice culture system"

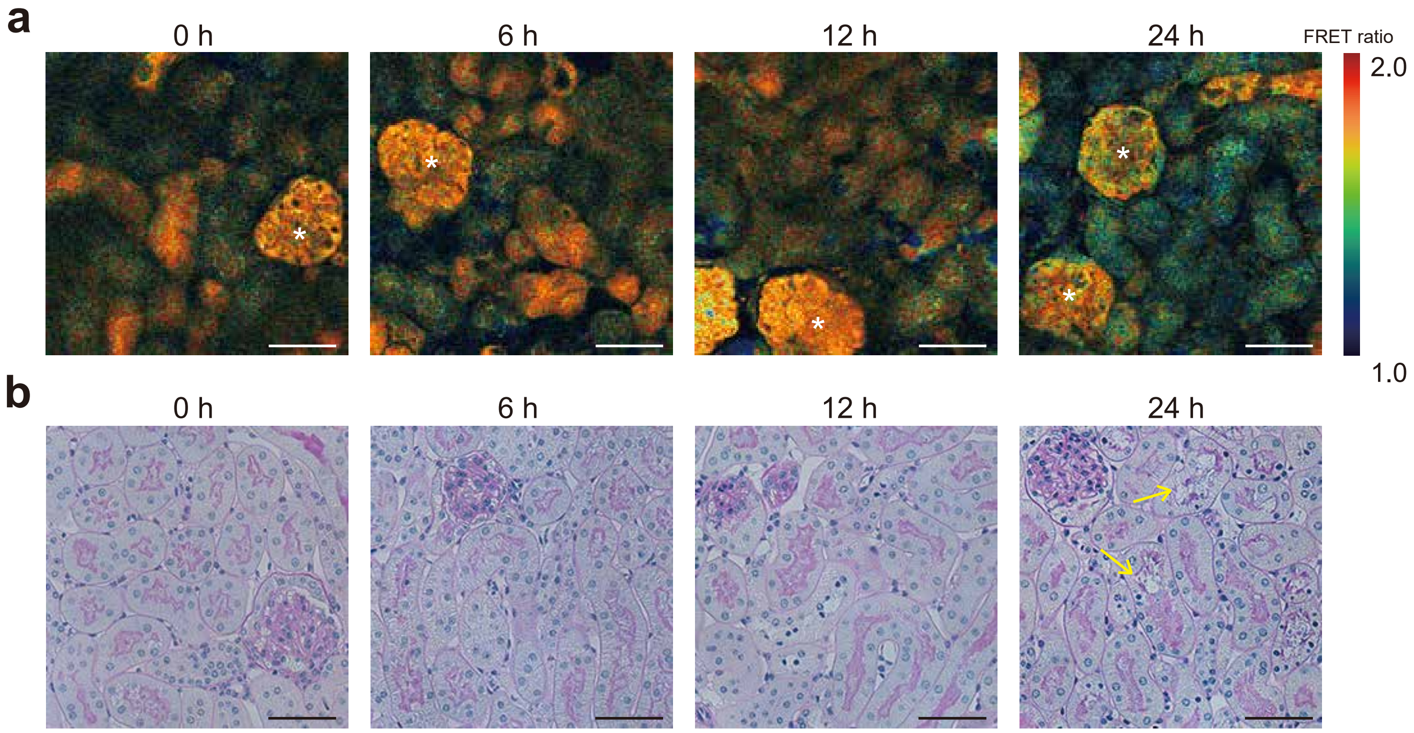


**Supplementary Figure S2. FRET ratio images and PAS staining of the kidney slices at longer time points after the kidney slice preparation.** (a) At 24 h after the kidney slice preparation, the ATP decline in some nephron segments including glomeruli (*) was observed. (b) PAS staining samples at 24 h after the kidney slices preparation showed loss of brush borders in PTs (arrows). Scale bars: 50 μm.

FRET, fluorescence resonance energy transfer; PAS, Periodic acid–Schiff; PTs, proximal tubules.
