## Supplementary Figure S3 for "Visualization of intracellular ATP dynamics in the whole kidney under pathophysiological conditions using the kidney slice culture system"

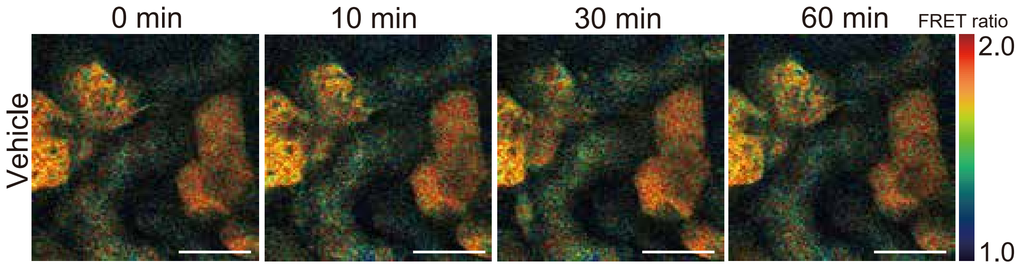


**Supplementary Figure S3. FRET ratio images after vehicle administration.** No apparent FRET ratio changes were observed in any segment. Scale bars: 50 μm.
