## Supplementary Figure S4 for "Visualization of intracellular ATP dynamics in the whole kidney under pathophysiological conditions using the kidney slice culture system"

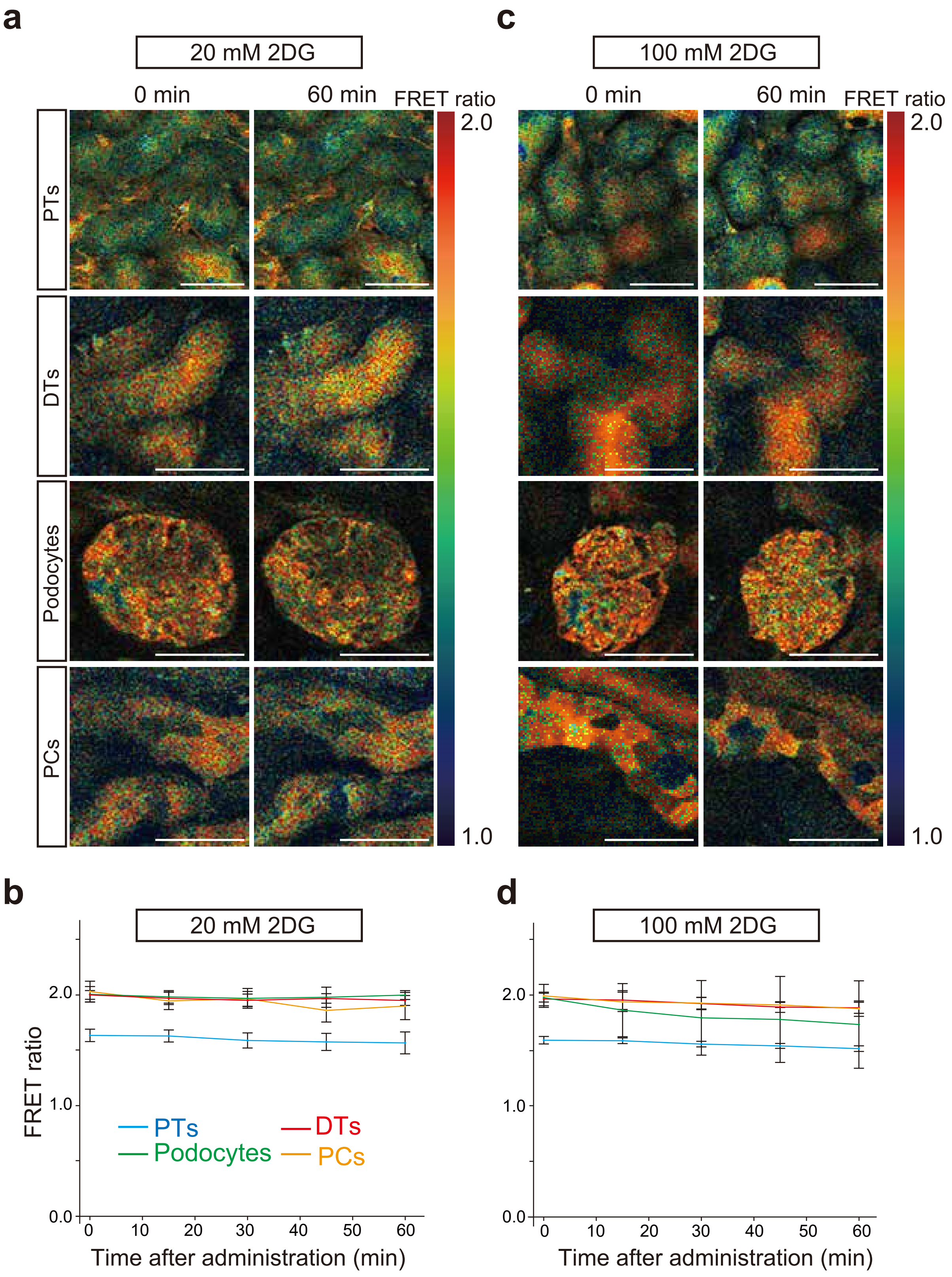


**Supplementary Figure S4. Intracellular ATP dynamics after 2DG administration.** (a, b) FRET ratio images and FRET ratio graphs in each nephron segment after 20 mM 2DG administration (*n* = 6 slices). (c, d) FRET ratio images and FRET ratio graphs in each nephron segment after 100 mM 2DG administration (*n* = 7 slices). Scale bars: 50 μm.
2DG, 2-deoxy-D-glucose; PTs, proximal tubules; DTs, distal tubules; PCs, principal cells.

.
