## Supplementary Figure S5 for "Visualization of intracellular ATP dynamics in the whole kidney under pathophysiological conditions using the kidney slice culture system"

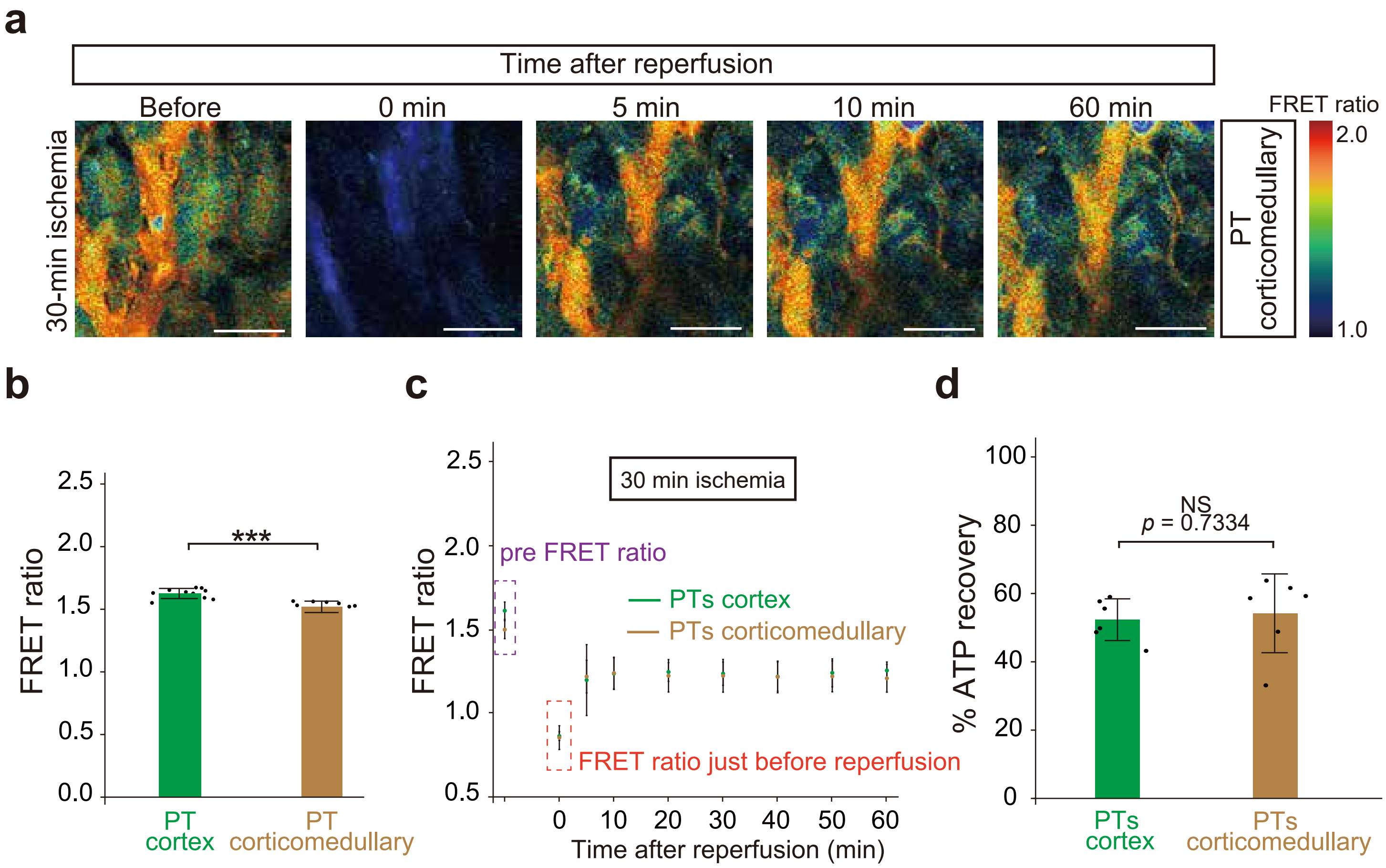


**Supplementary Figure S5. Intracellular ATP dynamics of the proximal tubules in the corticomedullary region during ischemia reperfusion injury.** (a) FRET ratio images of the proximal tubules (PTs) in the corticomedullary region during reperfusion after 30-min ischemia. (b) Pre FRET ratios (FRET/GFP) in PTs in the cortex and corticomedullary region ($n$ = 10 slices). The FRET ratio graphs of PTs in the cortex in Figure 1e are presented here again. (c) FRET ratio graphs of PTs in the cortex and corticomedullary region during reperfusion after 30-min ischemia ($n$ = 6 slices). FRET ratio graphs of PTs in the cortex after 30-min ischemia in Figure 3c are presented here again. (d) The % ATP recovery 60 min after reperfusion. Statistical significance was assessed using unpaired, two-tailed $t$-test. *** $p < 0.001$; NS, not significant. Scale bars: 50 μm.

FRET, fluorescence resonance energy transfer; PTs, proximal tubules.
