## Supplementary Figure S6 for "Visualization of intracellular ATP dynamics in the whole kidney under pathophysiological conditions using the kidney slice culture system"

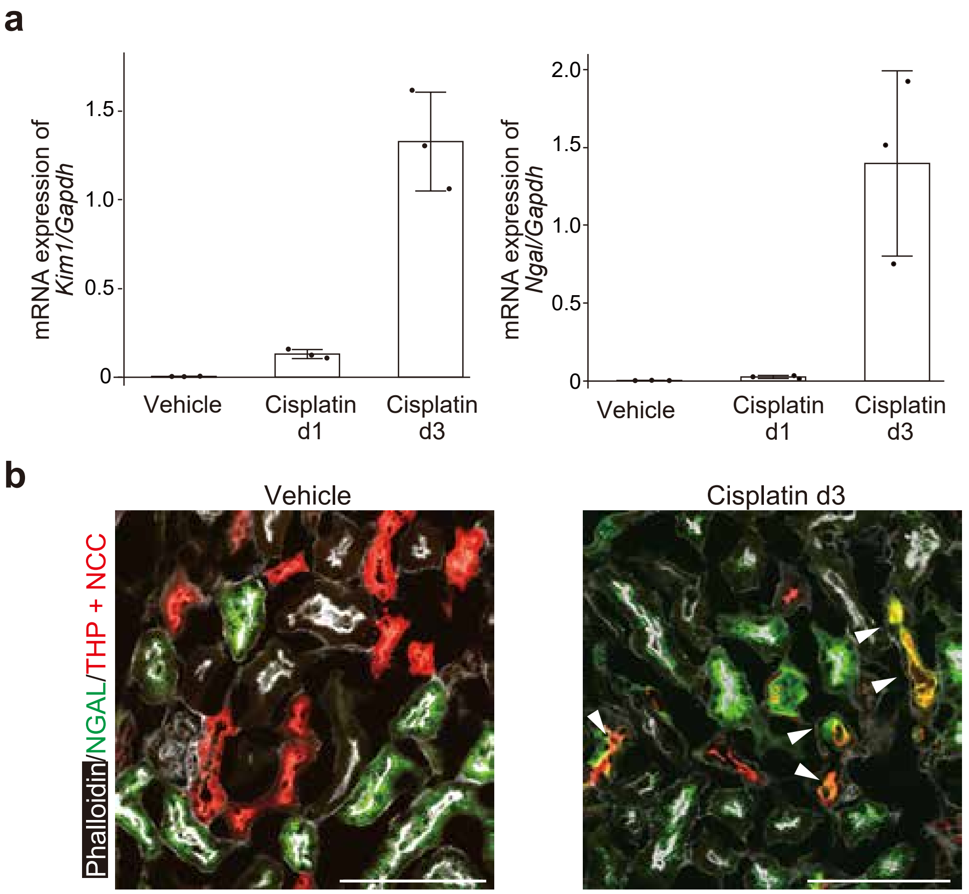


**Supplementary Figure S6. In addition to proximal tubule injury, cisplatin nephropathy causes distal tubule injury *in vivo.*** (a) Real-time PCR analysis showed elevated *Kim1* and *Ngal* mRNA expression levels in the kidneys of mice treated with cisplatin ($n$ = 3 mice per group). (b) Immunostaining analysis revealed NGAL staining colocalized with THP and NCC staining (arrowheads) in mice that were administered cisplatin, indicating distal tubule injuries. Positive NGAL staining in proximal tubules indicates reabsorption of NGAL from the urine. Scale bars: 100 μm. NGAL, neutrophil gelatinase-associated lipocalin; THP, Tamm–Horsfall protein; NCC, sodium chloride co-transporter.
