## Supplementary Figure S7 for "Visualization of intracellular ATP dynamics in the whole kidney under pathophysiological conditions using the kidney slice culture system"

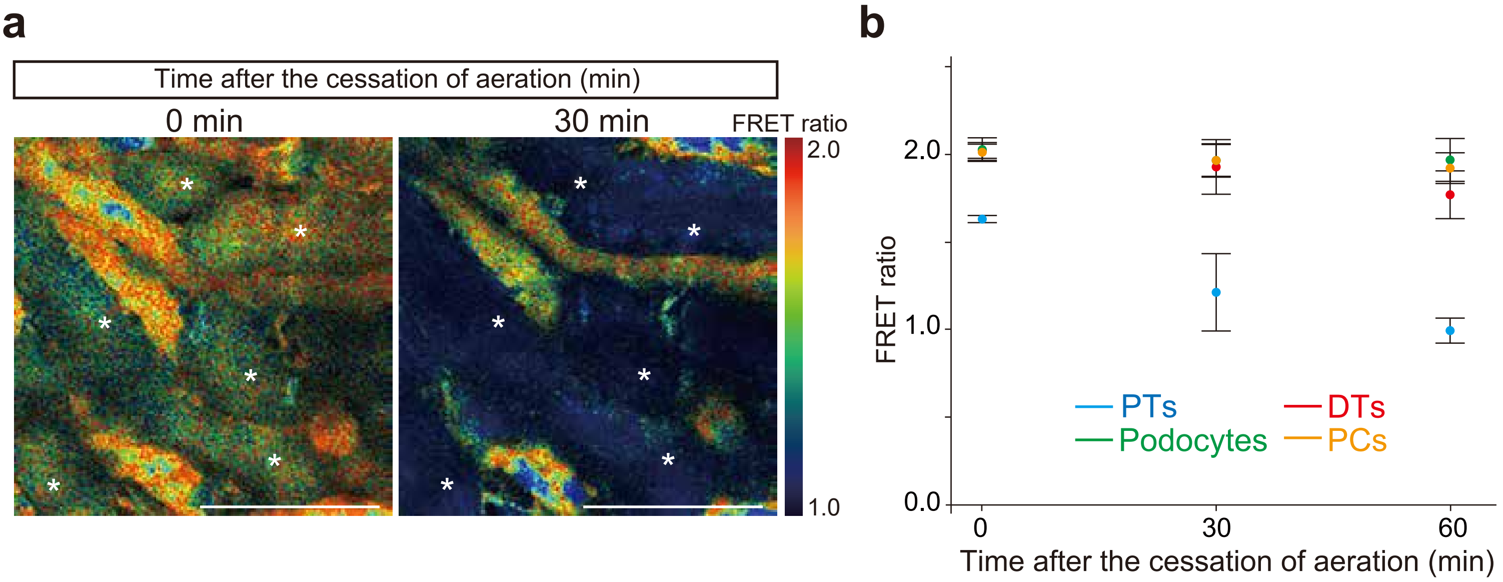


**Supplementary Figure S7. Intracellular ATP dynamics after the cessation of aeration.** (a) ATP levels in proximal tubules (*) decreased without aeration. (b) FRET ratio graphs in each nephron segment after the cessation of aeration (*n* = 5 slices). Scale bars: 100 μm. PTs, proximal tubules; DTs, distal tubules; PCs, principal cells.
