## Supplementary Table S1 for "Visualization of intracellular ATP dynamics in the whole kidney under pathophysiological conditions using the kidney slice culture system"

Sequences of primers used for real-time PCR

| Gene | Sequence (5’ − 3’) | |
|---|---|---|
| | Forward | Reverse |
| *Gapdh* | CCAGAACATCATCCCTGCATC | CCTGCTTCACCACCTTCTTGA |
| *Kim1* | TCTATGTTGGCATCTGCATCG | GAAGGCAACCACGCTTAGAGA |
| *Ngal* | AGGGCTGGCCAGTTCACTCT | CCATGGCGAACTGGTTGTAGT |
| *OCT2* | TGGCATCGTCACACCTTTCCT | CCAGCAACAAGGCCAACCAC |
