## Supplementary Methods for "Visualization of intracellular ATP dynamics in the whole kidney under pathophysiological conditions using the kidney slice culture system"

### *Animals*

GO-ATeam2 mice, which systemically express a fluorescence resonance energy transfer (FRET)-based ATP biosensor, were generated in our laboratory.[1,2] C57BL/6J mice were purchased from Japan SLC Inc (Shizuoka, Japan). The mice were housed in a specific pathogen-free facility, and received a routine diet; 10–13-week-old male mice were used in this study. All animal experiments were approved by the Animal Research Committee of the Graduate School of Kyoto University (No. MedKyo22183) and performed in accordance with the Guide for the Care and Use of Laboratory Animals (NIH).

### *Microscopy and image processing*

*Fluorescence stereomicroscopy*: The kidney slices were exposed to an excitation light (ET470/40) using a fluorescence stereomicroscope (Leica M165FC; Leica Microsystems). The fluorescent light was separated into green fluorescent protein (GFP) (D515/30m, Chroma Technology, VT, USA) and Kusabira orange fluorescent protein (OFP) (D575/40m, Chroma Technology) using a Dual-View (DM540; Nippon Roper, Tokyo, Japan), and the images were captured using a cMOS camera (ORCA-Flash4.0; Hamamatsu Photonics, Hamamatsu, Japan) to obtain dual-images simultaneously.

*Two-photon microscopy*: An FV1200MPE-BX61WI upright microscope equipped with a ×25/1.05NA water-immersion objective lens (XLPLN25XW-MP; Olympus, Tokyo, Japan) and an InSight DeepSee Ultrafast Laser (Spectra-Physics, Mountain View, CA, USA) were used. The excitation wavelengths for GFP and tetramethyl rhodamine methyl ester (TMRM) were 930 and 860 nm, respectively. We used an infrared (IR)-cut filter (BA750RXD), two dichroic mirrors (DM505 and DM690), and two emission filters (BA495-540 [Olympus] for GFP, and BA562-596 [Olympus] for Kusabira OFP and TMRM, respectively). Images were analyzed using the MetaMorph software (Universal Imaging, West Chester, PA, USA). All images were captured at approximately 10 μm from the cutting surface for analysis.

### *Mice treatment*

We administered 15 mg/kg cisplatin or vehicle intraperitoneally to 10–13-week-old GO-ATeam2 male mice and observed ATP dynamics in the kidney 3 days after the administration. As previously reported,[1] mice were anesthetized with 2% isoflurane inhalation, and the left kidney was exteriorized through a small incision. TMRM analysis was used to evaluate the mitochondrial membrane potential of the tubules after cisplatin or vehicle administration to C57BL/6J mice; 100 μl/30 g body weight of 0.3 mM TMRM was injected intravenously immediately before observation.

### *Renal histological analysis*

Kidney samples were fixed in Carnoy’ s solution, embedded in paraffin, sectioned (2.0 μm), and stained using periodic acid–Schiff (PAS) staining. All PAS-stained samples were analyzed using a Zeiss Axio Imager A2 microscope and Zeiss Axio (Carl Zeiss, Oberkochen, Germany) Vision 4.8 software.

***Renal immunostaining***

Kidney samples were fixed in 4% paraformaldehyde, incubated overnight in 20% sucrose in phosphate-buffered saline (PBS), and then incubated overnight in 30% sucrose in PBS at 4℃. Optical Cutting Temperature Compound (OCT)-embedded kidneys were cryosectioned into 6.0-μm sections. Kidney slices were immunostained directly after fixation. The following primary antibodies were used: antibodies against thiazide-sensitive NaCl cotransporter (NCC) (catalog no. AB3553; Millipore Corporation, Temecula, CA, USA), aquaporin 2 (AQP2) (catalog no. 178612; Calbiochem, San Diego, CA, USA), Tamm-Horsfall protein (THP) (catalog no. MAB5175; R&D systems, Minneapolis, MN, USA), nephrin (catalog no. AF3159; R&D Systems), GFP (catalog no. ab13970; Abcam, Cambridge, UK), Kim1 (catalog no. 14-5861; eBioscience, San Diego, CA, USA), and neutrophil gelatinase-associated lipocalin (NGAL) (catalog no. AF1857; R&D Systems).
Alexa-conjugated phalloidin (catalog nos. A22283, A22287; Invitrogen, Carlsbad, CA, USA) was utilized to visualize the brush borders of PTs.

***Quantification of mRNA expression using real-time polymerase chain reaction***

RNA extraction and real-time polymerase chain reaction (PCR) were performed as described previously.[3] Specific primers were designed using Primer-BLAST software. The sequences of the primers used for real-time PCR are listed in Supplementary Table S1

***Immunoblotting analysis***

Fresh snap-frozen kidney samples were homogenized in a lysis buffer containing protease and phosphatase inhibitors. The protein concentrations were determined using a BCA kit (TaKaRa Bio Inc., Shiga, Japan).
The samples were then subjected to sodium dodecyl sulfate polyacrylamide gel electrophoresis (SDS-PAGE) and immunoblot analyses. The following primary antibodies were used: organic cation transporter 2 (OCT2) (catalog no. OCT2 1-A; Alpha Diagnostic, San Antonio, TX, USA) and glyceraldehyde-3-phosphate (GAPDH) (catalog no. 10R-G109A; Fitzgerald, North Acton, MA, USA)

***Measurement of cisplatin concentration in the tissues***

The amount of cisplatin taken up by the kidney slices was determined using inductively coupled plasma-mass spectrometry (ICP-MS), as described elsewhere.[4,5] The kidney slices were incubated for 2 h in the following three groups: control; 1.0 mM cisplatin; and 1.0 mM cisplatin + 1.0 mM cimetidine. The same concentration of cimetidine was used in the other experiments. The excised kidney slices were washed rapidly three times using ice-cold buffer, weighed, snap-frozen, and stored at −80℃ until analysis. Kidney slices were minced in normal saline (9 μl/mg tissue); 60% HNO3 (40 μl/mg tissue) was added to lyse the tissues during incubation at room temperature overnight, followed by 95°C for 1 h. The lysates were further diluted 100-fold with 5% HNO3, and the cisplatin concentration was determined using ICP-MS (Agilent7700, Agilent Technologies, Santa Clara, CA, USA).

***Reagents***

Oligomycin A (catalog no. 75351; Sigma-Aldrich, St. Louis, MO, USA), 2-deoxy-D-glucose (2DG) (catalog no. 046-06483; Wako, Osaka, Japan), phloretin (catalog no. P7912; Sigma-Aldrich), cisplatin (catalog no. P4394; Sigma-Aldrich), cimetidine (catalog no. C4522; Sigma-Aldrich), and TMRM (catalog no. T668; Invitrogen) were purchased from their respective suppliers. Mitochonic acid 5 (MA-5) was synthesized chemically, as previously reported.[6]

***Statistics***

Results are presented as the mean ± standard deviation (SD) (Figure 1e, Figure 2, Figure 3, c–e, Figure 5c, Figure 6c, Figure 7, a, c, g and h, Figure 8, b and c; Supplementary Figure S4, b and d, Supplementary Figure S5, b–d, Supplementary Figure S6a, Supplementary Figure S7b), or as the median and interquartile range (IQR) (Figure 4b, Figure 5b, Figure 7e). Significant differences were determined using unpaired, two-tailed *t*-tests for comparisons between two groups (Figure 3e [between 30 min IR PTs and DTs], Figure 5, b and c, Figure 7a, Figure 8c; Supplementary Figure S5, b and d); one-way analysis of variance (ANOVA) with Tukey–Kramer *post hoc* tests for comparisons among more than two groups (Figure 1e, Figure 2, e and f, Figure 7, c, e and h); and a nonparametric test for trend (Cuzick’ s test for trend) across 15-, 30-, and 60-min IR groups (Figure 3e, Figure 4b). Statistical significance was defined as $p < 0.05$ using JMP Pro ver.15.2.0 software (SAS Institute, Cary, NC, USA) and R ver.4.1.2.
